# Deficiency in hyaluronan synthase 3 attenuates ruptures in a murine model of abdominal aortic aneurysms by reduced aortic monocyte infiltration

**DOI:** 10.1101/2022.12.01.518480

**Authors:** Tatsiana Suvorava, Fedor Brack, Janet Kaczur, Patrick Petzsch, Karl Köhrer, Christine Quast, Nobert Gerdes, Katharina Voigt, Martina Krüger, Jens W. Fischer, Alexander Brückner, Bernd K. Fleischmann, Daniela Wenzel, Laura-Maria A. Zimmermann, Gerhard Sengle, Ulrich Flögel, Maria Grandoch

## Abstract

Abdominal aortic aneurysms (AAA) are a common vascular disorder with a high mortality due to the prevalence of aortic ruptures. The underlying pathomechanisms are complex and involve immune cell infiltration and degradation of the vascular extracellular matrix (ECM). Hyaluronan (HA), synthesized at the plasma membrane by three HA synthase isoenzymes (HAS1-3), is not only a major constituent of the ECM but also known to directly affect the phenotype of vascular smooth muscle cells as well as immunological responses. Specifically, the HAS3 isoenzyme has been reported to play a major role in various inflammatory conditions. Therefore, the aim of the present study was to elucidate the role of HAS3-derived HA in the pathogenesis of abdominal aortic aneurysm. To this end, we used a murine model of Angiotensin II (AngII)-induced abdominal aortic aneurysms and dissections (AAAs/AADs) and could demonstrate that genetic depletion of *Has3* improves survival in *Apoe/Has3* double deficient (*Apoe/Has3*-DKO) mice via the reduced occurrence of aortic ruptures. Mechanistically, fewer elastica breaks were observed in *Apoe/Has3*-DKO mice compared to *Apoe*-KO littermates. This was associated with a decreased infiltration of myeloid immune cells into the vessel wall of *Has3*-deficient mice while in parallel elevated numbers of circulating leukocytes were detected. RNA seq analysis from aortic tissue pointed towards a disturbed endothelial-myeloid cell communication as a cause for the diminished recruitment of immune cells to the aortic wall. While endothelial cells were unaffected, upregulation of adhesion receptors as well as the HA receptor CD44, known to mediate leukocyte adhesion to the endothelium, was blunted in monocytes from *Apoe/Has3*-DKO mice in response to AngII treatment. These findings underline the pivotal detrimental role of monocyte’s HAS3-dependent pericellular HA matrix for an exaggerated immune cell recruitment to inflammatory foci giving here rise for an increased incidence of ruptured aortic aneurysms.

## Introduction

Aortic aneurysms are a common vascular disease associated with a high risk of morbidity and mortality (1, 2) The pathophysiology of abdominal aortic aneurysm (AAA) formation, a dilatation mostly occurring between the renal arteries and the bifurcation, includes diverse genetic and environmental risk factors such as age, gender and smoking (1, 2). During disease progression, the aortic wall becomes thinner and is prone to aortic dissections and ruptures as the diameter of AAA increases. Aortic dissections are described as a tear in the intima layer of the aorta, which then can penetrate the media and lead to a false lumen formation with or without communication to the true lumen of the aorta (3). A progressive separation of the aortic wall layers can finally lead to an aortic rupture, which then quickly results in exsanguination and followed by death.

The process of aneurysm formation in the abdominal aorta is characterized by a continuous degradation of the extracellular matrix (ECM), smooth muscle cell (SMCs) apoptosis as well as phenotypic switching (4, 5) and the infiltration of immune cells resulting in elastic fibre fragmentation (6, 7). The vascular hyaluronan (HA)-rich matrix is crucially involved in all of the above-mentioned processes, playing a fundamental role in the regulation of vascular function (8) and in the maintenance of aortic homeostasis (9). HA, a glycosaminoglycan synthesized by three HA synthase isoenzymes (HAS1-3), is not only located in all layers of the vascular wall (glycocalyx, media and adventitia) but also confers phenotypic modulations of mesenchymal cells and leukocytes, creating an adhesive matrix for monocytes and T cells (3, 7).

Inflammation is critical step in AAA pathogenesis and progression and is driven particularly by the recruitment of leukocytes to the vascular wall (10). The mononuclear macrophages, T- and B-lymphocytes are the predominant cell populations infiltrating the aortic wall in aortic aneurysm. Clinical studies revealed substantial modifications in peripheral blood monocyte subsets and phenotypes, suggesting the involvement of monocytes in regulating vascular inflammation and remodelling during the development of the disease (11). The numbers and phenotypes of monocytes were also significantly altered in abdominal aortic dissection (AAD) patients (12), suggesting that monocytes may play an important role in promoting the development AAD

In the acute phase proinflammatory Ly6C^high^ monocytes are recruited from the blood to the tissue via different chemokine receptors such as Ccr2, Ccr5, and Cx3cr1. Within the wall, monocytes differentiate to macrophages which in turn further enhance the release of proinflammatory cytokines/chemokines and matrix metalloproteinases. These inflammatory mediators degrade the ECM collagen and elastin followed by progressive luminal dilation. Collagen and elastin are mainly breakdown by a family of endopeptidases MMPs, with MMP9 expressed by infiltrating macrophages and SMC in the aneurysm lesions being crucial for AAA formation (13). Further, the differentiation to M1 macrophages resulted in the apoptosis of SMCs (14). Likewise, neovascularization of the medial layer which is a consistent feature of established AAA and aortic rupture (15) and reported to be caused by macrophages (3), can promote the separation and destruction of the aortic wall (16). In addition, local inflammation further causes ischemia, degeneration, and necrosis of aortic smooth muscle cells (11, 17) and changes of the flow dynamics may eventually lead to rupture of AAA.

In such inflammatory conditions, HA as one major ECM component is shaping macrophage as well as T cell responses. Thus, HA fragments can activate macrophages and increase the expression of inflammation-related cytokines (18). Further, the interaction between HA and HA receptors, such as CD44 and receptor of HA-mediated motility (RHAMM), modulate the adhesion, migration, and proliferation of SMCs (9, 19). In particular the HAS3 isoenzyme has been investigated in multiple cardiovascular pathologies and shown to be critically involved in these processes (19-23).

Therefore, this study aimed to elucidate the role of HA and specifically HAS3 in the the development of AAA and characterize the underlying HAS3-mediated mechanisms.

## Materials and Methods

### Mice

All experimental procedures and animal care were approved by the local animal experimentation ethics committee (LANUV; State Agency for Nature, Environment and Consumer Protection, file number 81-02.04.2018.A222), and performed in accordance with the guidelines of German Animal Welfare Law. Male mice double-deficient for apolipoprotein E (*Apoe*) and hyaluronan synthase 3 (Has3) (*Apoe/Has3*-DKO) and respective control mice (*Apoe*-KO) were used in this study. Has3-deficient mice were generated by genOway (Lyon Cedex, France) as described by Kiene et al. (19). For *Apoe/Has3*-DKO mice Has3-deficient mice were crossbred with *Apo*e-KO mice (Taconic, Hudson, NY, USA).

The animals were housed in a temperature-controlled room with a 12-hour light/dark cycle and had free access to water and food. All mice were fed a western-type diet (WD) containing 21% fat and 0.15% cholesterol (S8200-E010; Ssniff Spezialdiäten GmbH, Soest, Germany) starting from the day of surgery. Mice were expected daily and excluded from the experiment when certain criteria of suffering were observed. These included weight loss greater or equal than 20 % of body weight, cessation of food and water ingestion or lack of voluntary movement. At the end of the experiments, mice were sacrificed by CO_2_ inhalation, blood was obtained by cardiac puncture and organs were harvested for further analysis.

### AngII Infusion model of AAA

Eight to twelve weeks old Apoe-KO and DKO mice were infused with AngII or saline using osmotic minipumps (model 1004, Alzet, Cupertino, USA) at 1000 ng/kg/min as described by Trachet et al. (24) for 3, 7 and 28 days. The surgery was performed under anesthesia with ketamine (100mg/ml)/xylazine (5mg/kg i.p.). For pain relief, all animals received carprofen (5mg/kg body weight s.c.) immediately after surgery and on the next morning. Any mouse that died prior to the study endpoint underwent autopsy to determine cause of death.

### Porcine pancreatic elastase perfusion model

Porcine pancreatic elastase (PPE) perfusion model was used to induce abdominal aortic aneurysms as previously described by Pyo et al. (25). Mice were anesthetized with 2-3% isoflurane and median laparotomy was performed. After toe reflex was no longer present laparotomy was performed and the proximal and distal infrarenal aorta was isolated and temporarily ligated. The aorta was punctured, a catheter was carefully inserted and the infrarenal part was perfused with sterile isotonic saline containing type I Porcine Pancreatic Elastase (2.5-3 U ml, Sigma-Aldrich) for 5 min at 120 mm Hg. Elastase concentrations ranged from 2.5 to 3 U/ml depending on the LOT number, as different LOT numbers require different concentrations to trigger the same incidence of aneurysm. For postoperative pain mice received buprenorphine (TEMGESIC®, 0.1 mg/kg body weight s.c.) 30 min prior the surgery and carprofen 5 mg/kg/body weight s.c. for at least 3 days).)

### Blood Pressure Telemetry

Under ketamine/xylazine anesthesia mice were surgically implanted with a microminiaturized electronic pressure-sensing catheters (PA-C10, Data Sciences International (DSI), St. Paul, MN, USA) into the left common carotid artery. Digitized hemodynamic data were continuously sensed, processed and transmitted via radiofrequency signals to a nearby receiver (acquisition period: 10 min every hour). For pain relief buprenorphine (TEMGESIC®, 0.1 mg/kg body weight s.c.) was applied 30 min prior the DSI transmitter implantation and every 4 hours (s.c.) and in drinking water (0.009 mg/ml) for at least 3 days after surgery. Mice were allowed a 7-day post-surgery stabilization period before starting the acquisition of hemodynamic data. After 3 days of baseline measurement, AngII pumps were implanted as described above and blood pressure measurements were collected continuously with sampling every 20 min for 10 s intervals.

### Magnetic Resonance Imaging

Experiments were performed at a vertical 9.4 T Bruker AVANCE^III^ Wide Bore NMR spectrometer (Bruker) operating at frequencies of 400.21 MHz for ^1^H and 376.54 MHz for ^19^F measurements using microimaging units as described previously (26). Mice were anesthetized with 1.5% isoflurane and were kept at 37°C during the measurements. For gated MRI acquisitions, the front-paws and the left hind-paw were attached to ECG electrodes (Klear-Trace) and respiration was monitored by means of a pneumatic pillow positioned at the animal’s back. Vital functions were acquired by a M1025 system (SA Instruments) and used to synchronize data acquisition with cardiac and respiratory motion. Data were acquired using a 25-mm quadrature resonator tuneable to ^1^H and ^19^F. After acquisition of the morphological ^1^H images, the resonator was tuned to ^19^F and anatomically matching ^19^F images were recorded. The reference power and the receiver gain were kept constant between the measurements to ensure comparability of the ^19^F scans.

### In Vivo ^1^H MRI

To conduct ^1^H/^19^F MRI, perfluorocarbon nanoemulsions (PFCs) were prepared as described previously (26) and injected intravenously (3 mmol/kg/BW) via the tail vein in anesthetized (1.5% isoflurane) mice. To visualize the anatomy of the region of interest, ^1^H MR reference images from the abdomen were acquired using a rapid acquisition and relaxation enhancement sequence [RARE; field of view (FOV) = 2.56 × 2.56 cm2, matrix = 256 × 256, 0.1 × 0.1 mm2 in plane resolution, 1 mm slice thickness (ST); repetition time (TR) = 2,500 ms; RARE factor = 16, 6 averages (NA), acquisition time (TAcq) = ∼5 min] as described previously (27). ^1^H MR time-of-flight angiography to visualize dilatation or narrowing of the aorta via its blood flow pattern was carried out by a ^1^H fast low angle shot (FLASH) 2D flow compensated sequence; FOV = 2.56 × 2.56 cm2, matrix = 256 × 256, 0.1 × 0.1 mm2 in plane resolution, ST = 0.5 mm; 0.25 mm overlap, 100 slices; TR = 10 ms; NA = 6; TAcq = 4 min). Aortic bright blood cine movies were acquired in sagittal orientation adapted to the anatomical course of the vessel using an ECG- and respiratory-gated segmented fast gradient echo cine sequence with steady state precession (FISP). A flip angle (FA) of 15°, echo time (TE) of 1.23 ms, and a TR of about 6–8 ms (depending on the heart rate) were used to acquire 16 frames per heart cycle from a FOV of 30 × 22 mm2, matrix = 256 × 192, ST = 1 mm, NA = 3, TAcq per slice for one cine loop ∼2.5 min. From the same FOV, corresponding black blood cine movies were recorded utilizing an ECG- and respiratory-gated FLASH sequence with black blood preparation (blood inversion time ∼100 ms and TR ∼8 ms depending on the heart rate, TE = 2.28 ms, matrix = 256 × 192, ST = 1 mm, 16 frames, NA = 4, TAcq ∼5 min.

### In vivo ^1^H/^19^F MRI

Anatomically matching 19F images were recorded from the same FOV with a ^19^F RARE sequence (matrix = 64 × 64, 0.4 × 0.4 mm2 in plane resolution, ST = 1 mm, TR = 4,000 ms, RARE factor = 32, NA = 25, and TAcq = 34 min) as described (27).

### Quantification of Aneurysm *in vivo* by Ultrasound

Mice were monitored for aortic aneurysm formation at baseline and three days, one week, two weeks, three weeks and four weeks after implantation of minipumps with AngII or PPE perfusion using a Vevo 3100 high-resolution *in vivo* micro-imaging system and a 20-46 MHz - transducer (MX400) (VisualSonics Inc., FUJIFILM, Toronto, Canada). In the abdominal aorta ultrasound images were captured in long-axis view, short axis view was used to measure lumen area as maximum transverse dimension orthogonal to the vessel axis. In the thoracic aorta, the right parasternal long-axis view was used for ascending aortic imaging. All the measurements were performed at mid-systole, when the aorta is maximally and visually dilated. Color Doppler helped to visualize the blood flow pattern and aortic lumen in AAA. Pulsed wave (PW) Doppler was used to measure the velocity of blood either in suprarenal aorta between celiac artery and superior mesenteric artery (AngII model) or in infrarenal aorta (PPE model) model or the largest portion of aneurysm.

Ultrasound images were analyzed using VEVO 3100 software (VevoLAB, FUJIFILM, Visual Sonics Toronto, Canada). Longitudinal und short axis 2D B-modes and PW Doppler were used to assess maximal systolic and diastolic diameters, centerline blood velocity, visualize true und false lumen and thrombus within the dissection. Dissecting AA (AAD) were determined by their increase in diameter, changes in blood flow velocity and a presence of a dissection and flap in AAD. A blinded single investigator performed evaluations at each time point for the study groups.

### Quantification of elastin fragmentation

After reperfusion with PFA (%) mice were sacrificed, aortas were rinsed with PBS (Invitrogen, Carlsbad, CA, USA) then subsequently fixed in 4% formaldehyde solution (Histofix 4%, Roth, Karlsruhe, Germany). After 24 h aortas were transferred into PBS, drained, embedded in paraffin and sliced in 4 μm sections. Cross sections of maximum diameter and adjacent sections intervals were used for conventional Verhoeff-Van Gieson staining to assess the elastin architecture of the aortic aneurysm. Degree of fragmentation of the elastic fibers was examined using the number of breaks pro µm^2^.

### Tropoelastin Staining

Tropoelastin staining was performed on paraffin-embedded aortas. For the dewaxing of tissue sections, the slides were rehydrated with a decreasing alcohol concentration series. This was done in 5 minutes incubation steps in the following order: xylene I, II, isopropanol I, II, 96% EtOH, 80% EtOH, 70% EtOH, and finally distilled H_2_O. PFA-fixed tissue preparations were subjected to epitope retrieval by boiling in an evaporation dish at 180 W for 15 min to avoid bubble formation with a citrate buffer at pH 6.0. The buffer level was checked every 5 min, and was refilled, whenever necessary. After boiling, the evaporation dish was cooled together with the slides for 20 min at RT. After a TBS washing step and a one-hour blocking step with 5 % BSA / 0.1% Triton X-100 in 1 × TBS, the sections were incubated at 4 °C overnight with the primary rabbit polyclonal anti-tropoelastin antibody solution (Abcam, ab21600) 1:200 in 0.5% BSA in TBS. After two TBS-Tween 0.1% washing steps and a TBS washing step, the secondary anti-rabbit antibody (InvitroGen Thermo Fischer, Carlsbad, CA, USA), 1:1000 in 0.5 % BSA in TBS was applied for 1h. Afterwards, the slides were washed with big volumes of TBS-Tween 0.1 % and TBS and were stored in TBS buffer in the cold.

### Second Harmonic Generation Microscopy

For collagen detection, 10 µm crossections of paraffin embedded aortas were dewaxed, and second harmonic generation microscopy (SHG) of aortic collagens was obtained by using a Leica TCS SP8 microscopy (Leica microsystems, Wetzlar, Germany) with a Chameleon Vision II laser. SHG and TPEF were excited at 810 nm and imaged with an IR Apo L25x/0.95 W objective lens.

### Microscopy

Microscope pictures of aortic histological sections were taken using the IMAGER M2 microscope (Carl Zeiss Microscopy GmbH, Oberkochen, Germany) and analyzed with the ZEN software (Carl Zeiss Microscopy GmbH, Oberkochen, Germany).

### Flow Cytometry

For the flow cytometric analysis, blood and aortas were harvested 7 and 28 days after the Ang II minipump implantation. After collecting blood samples by heart puncture erythrocytes were lysed with hypotonic ammonium chloride solution. Aortae were dissected and digested in a solution containing of 1200 U/ml collagenase II (Worthington Biochemicals, Lakewood, NJ, USA), 60 U/ml DNase (Sigma Aldrich) for 60 minutes at 37°C. After digestion, aortas were filtered through 70 µm cell strainers (Greiner BioOne, Kremsmünster, Austria). Single cell solution was centrifuged at 300 x g for 10 minutes at 4°C, and resuspended in Dulbecco’s Modified Eagle Medium (DMEM, Life Technologies™, Thermo Fischer Scientific, Waltham, MA, USA) supplemented with 1 % fetal calf serum (FCS, Gibco, NY, USA) and incubated at 37°C for 30 minutes. After second centrifugation step at 300 g for 10 minutes at 4°C, cell pellets were resuspended in PBS containing 2 mmol/l EDTA and 0.5 % bovine serum albumin. To avoid unspecific binding, isolated cells were incubated with a CD16/32 antibody, before staining with LIVE/DEAD Fixable Aqua Dead Cell Stain Kit. A complete list of antibodies including the respective clones, manufacturers and dilutions and definitions of immune cell populations is given in **Suppl. Table 1 and 2** BD Fortessa™ Flow Cytometer (BD Biosciences, Heidelberg, Germany) was used to perform flow cytometry. Data were analyzed with FlowJo 10.8.1 Analysis Software (Becton Dickinson & Company (BD), Franklin Lakes, New Jersey, USA). For determination of total cell concentrations Flow-12 Count™ Fluorospheres (Beckman Coulter Inc., Krefeld, Germany) was used.

### Isolation of Bone Marrow-derived monocytes and In Vitro Incubation with AngII

Bone marrow cells were isolated from femurs and tibias as described (28). Isolation of primary monocytes from the bone marrow of 10-12 week-old female Apoe-KO and Apoe/Has3-DKO mice was performed by using the Monocyte Isolation Kit, mouse (Miltenyi Biotec, Bergisch Gladbach, Germany) according to the manufacturer’s instructions (29). Monocytes were cultured in 5.5 mmol/L glucose DMEM supplemented with 10% FBS, 15 mmol/L HEPES, 100 U/ml penicillin and 100 μg/ml streptomycin (all reagents from Gibco Life Technologies, Paisley, UK) and incubated with AngII (10 ng/ml) or respective vehicle control for 6h. Subsequently, total RNA was isolated and gene expression analysis by qPCR was performed as described below.

### RNA Isolation from Aortic tissue and Monocytes and Quantitative Real-Time Polymerase Chain Reaction

RNA was isolated from *Apoe*-KO and *Apoe/Has3-*DKO aortae and primary bone marrow derived monocytes by using the RNeasy® Plus Universal Kit Mini 50 (Qiagen GmbH, Hilden, Germany). RNA concentration and purity (260 nm/ 280nm) were spectrophotometrically determined on a Nanodrop (Peglab, Erlangen, Germany). cDNA was synthesized using the QuantiTect® Reverse Transcription Kit (Qiagen, Hilden, Germany).

Quantitative real-time PCR (qPCR) was performed with Platinum SYBR Green qPCR SuperMix-UDG (Life Technologies, Eugene, OR, USA) on a StepOnePlus Real Time PCR System (Applied Biosystems). Samples were measured in duplicate and relative mRNA expression was calculated using the 2-ΔΔCt method with *Rn18s* as an internal control (h18). Primers used for qPCR are listed in **Suppl. Table 3**.

### RNA Isolation of Murine Endothelial Cells Derived from the Abdominal Part of the Aorta

The aortas of male *Apoe*-KO and *Apoe/Has3*-DKO mice (8 weeks) were removed and the abdominal aorta was isolated by cutting just below the diaphragm and just above the common iliac arteries. Vascular rings were cut open and ECs were isolated using an ice-cold metal rod (6 mm diameter) according to the “modified Häutchen method” described by Simmons et al. (30). Then, RNA from ECs was isolated using the RNeasy Plus micro kit according to the manufacturer’s instructions (Qiagen, Hilden, Germany). To assess RNA quality the RNA integrity number (RIN) was determined by a 2100 Bioanalyzer (Agilent Technologies, Santa Clara, CA, USA). Only samples with a RIN above 5.0 were processed further.

### RNA-Seq Analyses of Murine Aortas

DNase digested total RNA samples used for transcriptome analyses were quantified (Qubit RNA HS Assay, Thermo Fisher Scientific) and quality measured by capillary electrophoresis using the Fragment Analyzer and the ‘Total RNA Standard Sensitivity Assay’ (Agilent Technologies, Inc. Santa Clara, USA). All samples in this study showed high quality RNA Quality Numbers (RQN; mean = 8.7). The library preparation was performed according to the manufacturer’s protocol using the “VAHTS™ Universal RNA-Seq Library Prep Kit for Illumina® V6 with mRNA capture module”. Briefly, 500 ng total RNA were used for mRNA capturing, fragmentation, the synthesis of cDNA, adapter ligation and library amplification.

The bead purified library pool was sequenced on the NextSeq2000 system (Illumina Inc. San Diego, USA) with a read setup of PE 2×100 bp. The BCLConvert tool (v3.8.4) was used to convert the bcl files to fastq files as well for adapter trimming and demultiplexing.

### RNAseq Analysis of Murine Endothelial Cells Derived from the Abdominal Part of the Aorta

For library preparation, the Trio RNA-Seq Library Preparation kit (TECAN, Männedorf, Switzerland) was used. Five PCR cycles were applied for library amplification and libraries with an average fragment size of 308 bp were sequenced on a NextSeq 550 in paired-end mode (75 bp) at the Genomics & Transcriptomics Labor at the Heinrich Heine Universität Düsseldorf (Düsseldorf, Germany). For bioinformatic analysis, we used the Galaxy platform (Freiburg Galaxy Project (31)). RNA sequencing reads were mapped using RNA STAR (32) followed by counting reads per gene by using featureCounts (33).

As an additional quality control step the purity of ECs in the respective sample was determined by analysing expression levels of the classical EC marker genes Pecam1, vWF and Cdh5. The normalized counts of these 3 marker genes were added up for each sample and compared with EC marker expression in control adventitial samples from 3 animals. Only samples with EC marker expression of >2-fold of the mean EC marker expression in adventitial samples were included in the analysis.

In the remaining samples (n=5 for Apoe-KO and n=7 for Apoe/Has3-DKO), differentially expressed genes were identified by DESeq2 (34). For data visualization and cluster analysis heatmap2 (31) (Freiburg Galaxy Project was used, for which the gene expression was further normalized using a shifted log transformation [log10 (n+1)].

### Statistical Analysis

Data analyses on fastq files were conducted with CLC Genomics Workbench (version 22.0.2, QIAGEN, Venlo. NL). The reads of all probes were adapter trimmed (Illumina TruSeq) and quality trimmed (using the default parameters: bases below Q13 were trimmed from the end of the reads, ambiguous nucleotides maximal 2). Mapping was done against the Mus musculus (mm39; GRCm39.104) (Mai 17, 2021) genome sequence. After grouping of samples (two to nine biological replicates each) according to their respective experimental condition, the statistical differential expression was determined using the Differential Expression for RNA-Seq tool (version 2.7). The Resulting *P* values were corrected for multiple testing by FDR and Bonferroni-correction. A *P* value of ≤0.05 was considered significant. The Gene Set Enrichment Test (version 1.2) was done with default parameters and based on the GO term ‘biological process’ (M. musculus; December 16, 2021).

Other statistics were performed using the GraphPad Prism 9 Software (La Jolla, CA, USA). If not differently specified, the results are given as mean ± standard deviation (SD). For repeated measurements, data were analyzed by 1-way or 2-way ANOVA as appropriate followed by Bonferroni’s or Sidak’s post hoc tests as indicated. Where indicated, unpaired Student’s t-test was used to determine if two groups of data were significantly different. Normal distribution was tested by the D’Agostino-Pearson test. p < 0.05 was considered as statistically significant.

## Results

In the present study the role of HAS3-derived HA was investigated in a murine model of AngII-induced AAA formation and rupture in male *Apoe-* and *Apoe/Has3*-deficient mice fed a Western type diet.

### Improved survival in *Apoe/Has3*-DKO mice after 28 days of AngII infusion

In a first set of experiments, 58 male Apoe-KO mice and 47 *Apoe/Has3-*DKO mice were implanted with AngII pumps at a delivery rate of 1 µg/kg/min for 4 weeks. All mice were set on a WD on the day of pump implantation. Four weeks of treatment with AngII led to a slight reduction in body weight during the first week of the treatment **(Suppl. Fig 1A)** and induced similar degree of cardiac hypertrophy in both strains **(Suppl. Fig. 1B**,**C)**. Thirty-six of the 58 *Apoe*-KO mice died from aortic rupture while in *Apoe/Has3-*DKO group only 19 out of 47 mice **(Figure 1A,B)** (survival rate of 37.9% in *Apoe*-KO and 59.6% in *Apoe/Has3-*DKO accordingly). Post mortem analysis showed that ruptures at the aortic arch were nearly as prevalent as in the suprarenal abdominal segment in both, *Apoe-*KO and *Apoe/Has3-*DKO.

**Figure 1.**
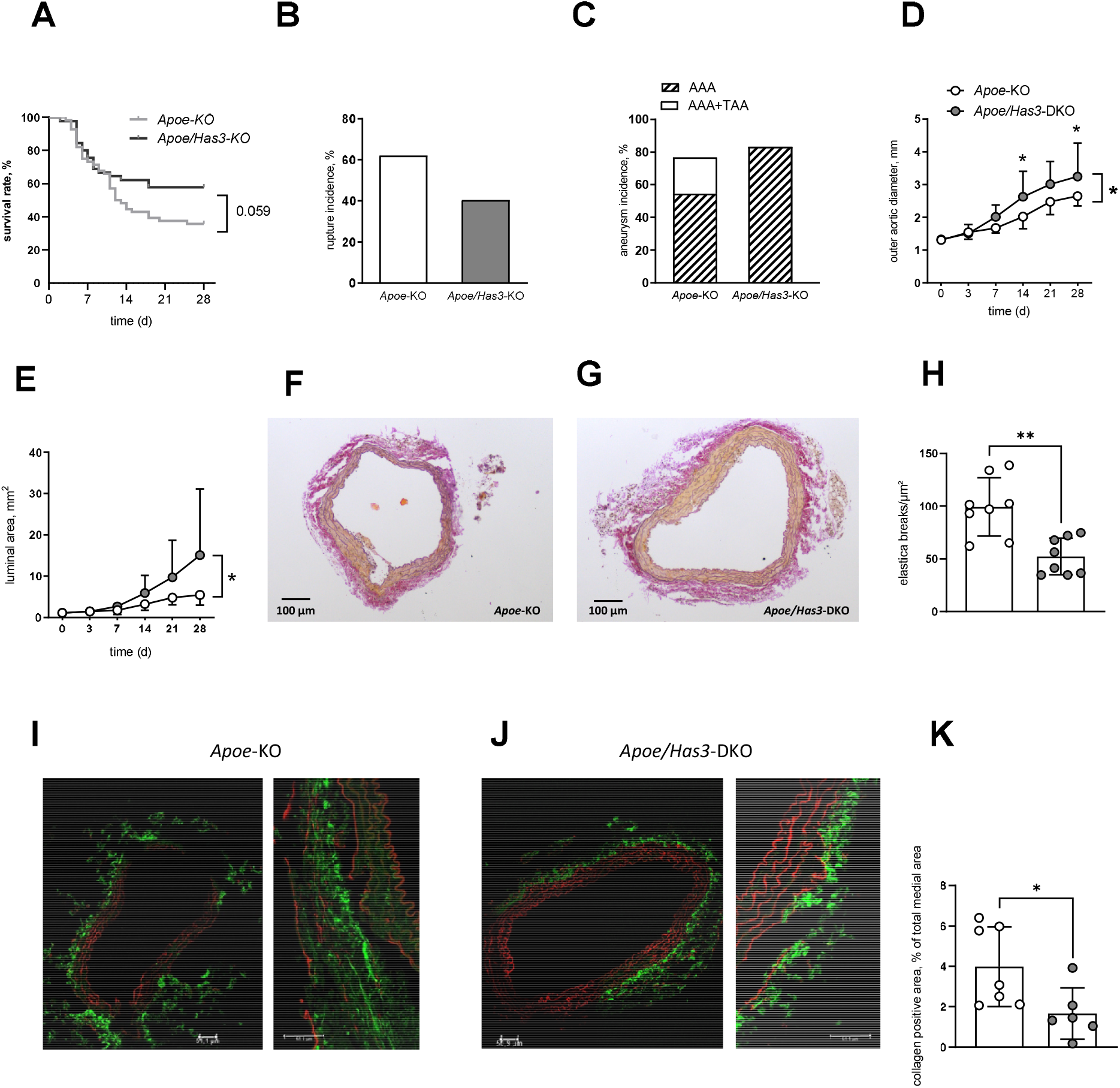
Hyaluronan synthase 3 (*Has3*)-deficiency protects against aortic rupture during 28 days of AngII infusion. **A**, Improved survival in *Apoe/Has3*-DKO mice. Kaplan-Meier curve represents the percentage survival in AngII-infused *Apoe*-KO (n=58) and *Apoe/Has3*-DKO (n=47), *P* = 0.059, Log-Rank (Mantel-Cox) test. **B**, Rupture incidence in *Apoe*-KO (n=58) and *Apoe/Has3*-DKO (n=47). **C**, The incidence of abdominal aneurysm formation (AAA) and thoracic aortic aneurysm (TAA) estimated as 1.5-fold increase of suprarenal or ascending aortic diameter in *Apoe*-KO and *Apoe/Has3*-DKO. **D**, Larger aortic external diameter and **E**, luminal area in AngII-treated *Apoe/Has3*-DKO (n=12) vs. *Apoe*-KO (n=11) measured by *in vivo* ultrasound,* *P* < 0.05, Two-way ANOVA followed by Sidak’s post hoc test. **F-G**, Fewer elastica breaks in aortic aneurysm tissue sections of *Apoe/Has*3-DKO. Representative Verhoeff-van Gieson staining of the elastic laminae in **F**, *Apoe*-KO and **G**, *Apoe/Has3*-DKO. **H**, Quantification of the number of ruptures of elastin fibers; n=8, ** *P* < 0.01, unpaired students *t*-test. **I - K**, Reduced collagen deposition in aortas of *Apoe/Has3*-DKO. Representative sections of the collagen deposition (green, label free) in the suprarenal aortic tissues of **I**, *Apoe*-KO and **J**, *Apoe/Has3*-DKO, **K**, Evaluation of the collagen positive area, n=7-6, * *P* < 0.05, unpaired students *t*-test. Data are presented as means ± SD.

### Increased size of aortic aneurysms in *Apoe/Has3*-deficient mice

In order to characterize aneurysm progression in more details, mice were monitored by ultrasound measurements longitudinally at baseline and on day 3, 7, 14, 21 and 28 of AngII infusion. The incidence of abdominal aneurysm formation (estimated as 1.5-fold increase in suprarenal diameter at day 28) was 54.5 %in *Apoe-*KO and 83.3 % in *Apoe/Has3*-DKO **(Figure 1C)**. There was a significant enlargement of abdominal suprarenal aorta after 4 weeks of AngII treatment in both *Apoe-*KO and *Apoe/Has3*-DKO, which surprisingly was more prominent in the abdominal aorta of *Apoe/Has3*-DKO: The transverse B-mode images showed an 100% increase of the aortic diameter (compared to baseline) in *Apoe*-KO and 150% in *Apoe/Has3*-DKO (**Figure 1D**). Likewise, AngII treatment induced a lumen extension, which was greater in *Apoe/Has3*-DKO (**Figure 1E, Suppl. Figure 1D,E)**. These aortic changes were preceded by a significant reduction of maximal flow velocity on day 3 of AngII infusion in both genotypes (**Suppl. Figure 1F**).

Taken together, these data demonstrate that mice deficient for *Has3* are characterized by a greater extension of the abdominal aorta during AngII infusion, which, however, does not lead to increased ruptures. Of note, the finding of larger aortic diameters might be misleading since more *Apoe*-KO mice have already died by rupture before day 28 thereby promoting a bias in the comparison of both groups.

### *Has3* deficiency limits aortic elastin breaks and collagen deposition

Since aneurysm rupture incidence was lower in *Apoe/Has3*-DKO mice we quantified elastin fragmentation in AAA tissue sections from AngII-infused *Apoe*-KO and *Apoe/Has3*-DKO using Verhoeff-van Giessen staining. Extensive destruction of elastin fibers within the aortic media, a hallmark of aneurysm pathology, was present in both strains on day 28 after AngII infusion. However, elastin fragmentation was significantly decreased and its structure and wall layering were better preserved in AAA sections of *Apoe/Has3*-DKO compared with *Apoe*-KO mice (**Figure 1F-H**).

These findings suggest that deficiency in *Has3* results in an improved survival and likely a more extensible abdominal aorta in mice at day 28 post-AngII infusion. As indicated above, this comparison is somewhat biased by a significantly higher survival rate in *Apoe/Has3-*DKO at this time point and thereby comparin*g Apoe/Has3*-DKO with fully formed AAA with the survived *Apoe*-KO which probably show the mildest phenotype. Therefore, we investigated both mice strains in a second model of AAA, the porcine elastase model. Nevertheless, also in this model, increased aortic diameters could be observed in *Apoe/Has3*-deficient mice compared to *Apoe*-KO mice (**Supplemental Figure 2**).

Next, to elucidate the mechanisms underlying the improved survival and fewer ruptures in *Apoe/Has3*-DKO, both genotypes were studied at an early stage of aneurysm development, i.e. within the first week of AngII infusion.

### Similar survival rate and AAA development but less elastic breaks and collagen deposition in the aorta of *Apoe/Has3-*DKO after one week of AngII treatment

One week of AngII infusion lead to a similar alterations in body weight (**Suppl. Figure 1A)** and cardiac hypertrophy in *Apoe*-KO and *Apoe/Has3*-DKO mice as described above (as evidenced by an increase of heart weight/body weight (**Suppl. Figure 3A,B)** and left ventricular mass to body weight ratios (**Suppl. Figure 3C**). However, transthoracic echocardiography revealed no differences in cardiac function in response to AngII infusion between *Apoe/Has3*-DKO and their controls during this time (**Suppl. Figure 3D-H**). Blood pressure and heart rate measured by radiotelemetry at baseline and during infusion of AngII were also not significantly different between *Apoe*-KO and *Apoe/Has3*-DKO neither during daytime (**Suppl. Figure 3I-L**) or nighttime (**Suppl. Figure 3M-P**).

The percentage of mice that survived after 1 week of the AngII infusion was similar in *Apoe* KO (71%) and *Apoe/Has3*-DKO (67%) mice (**Figure 2A**). AngII progressively increased abdominal suprarenal aortic diameter, and at days 7 of the Ang II infusion 26.9% of Apoe-KO and 27.7% of *Apoe/Has3*-DKO developed AAA (**Figure 2B**). At this early time point *Has3* deficiency had no effect on aortic diameter and intraluminal area (**Figure 2C-D**).

**Figure 2.**
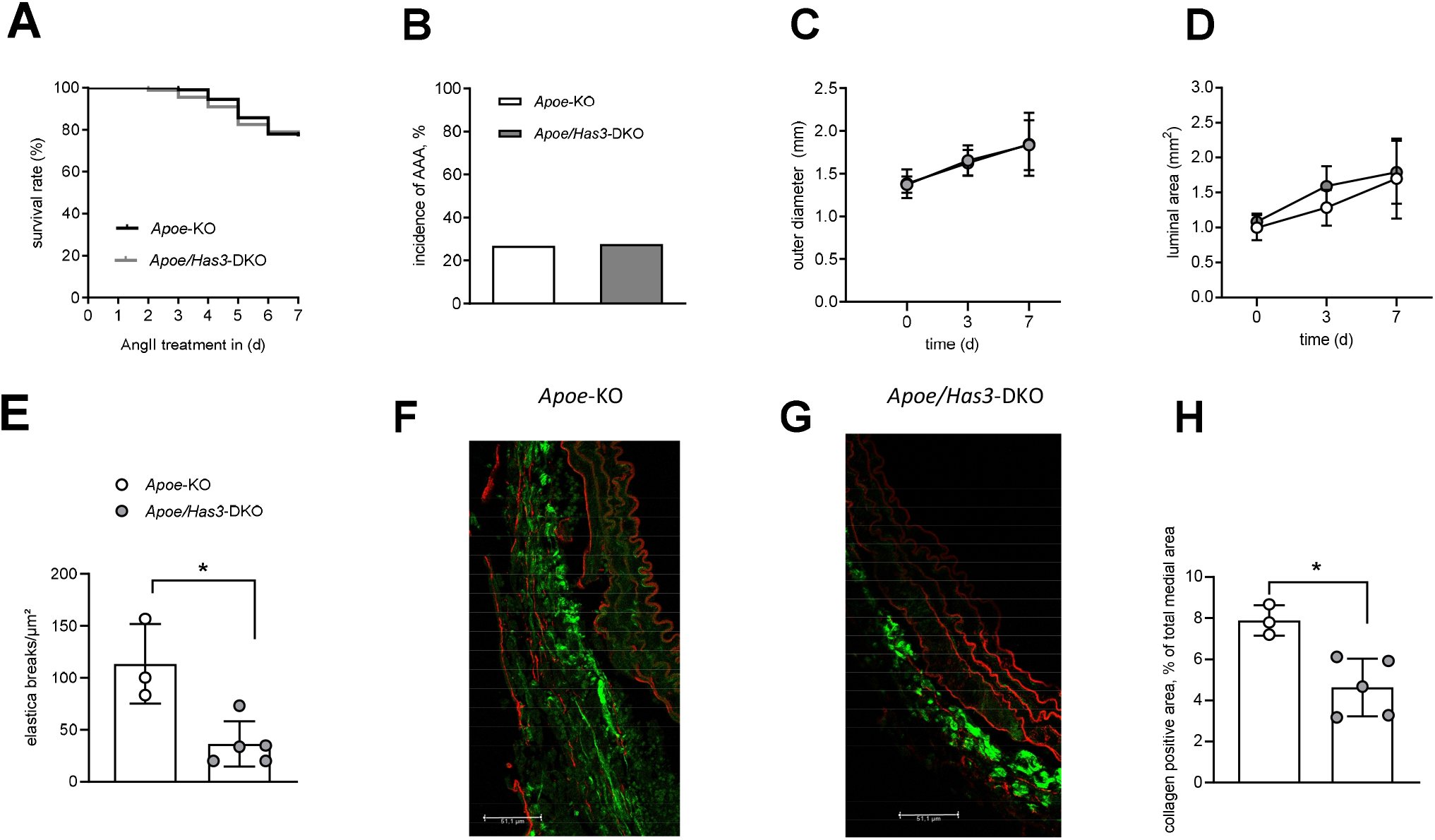
Similar survival rate and aortic aneurysm development in *Apoe*-KO and *Apoe/Has3*-DKO mice after one week of AngII treatment. **A**, The percentage of the mice that survived during 1 week of the AngII infusion was similar in *Apoe*-KO (n=95) and *Apoe/Has3*-DKO (n=90), p ≥ 0.05, Log-Rank (Mantel-Cox) test. **B**, Incidence of AAA was not different between *Apoe*-KO and *Apoe/Has3*-DKO after 1 week of the AngII infusion. **C**, Aortic suprarenal outer diameter and **D**, luminal area were not different between *Apoe*-KO (n=13) and *Apoe/Has3*-DKOs (n=11), *P* ≥ 0.05, Two-way ANOVA followed by Sidak’s post hoc test. **E**, Decreased number of elastica breaks in AAA from *Apoe/Has3*-DKO (n=8) vs *Apoe*-KO (n=8), * *P* < 0.05, unpaired students *t*-test. **F - H** Representative sections of the collagen deposition (green, label free) in the suprarenal aortic tissues of **F**, *Apoe*-KO and **G**, *Apoe/Has3*-DKO and **H**, Quantification of the collagen positive area, n=3-5, * *P* < 0.05, unpaired students *t*-test. Data are presented as means ± SD.

As observed also after 28 days of AngII treatment, number of elastic breaks was decreased in AAA sections from *Apoe/Has3*-DKO mice compared with *Apoe*-KO (**Figure 2E)**. Moreover, the collagen positive area was significantly decreased in AAA tissue sections from AngII-infused *Apoe/Has3-*DKO as compared to *Apoe*-KOs (**Figure 2F-H**) suggesting diminished collagen deposition in *Has3* deficiency after 7 days of AngII infusion.

### *Has3* deficiency attenuates the inflammatory cell infiltration into the aortic wall

The activation of inflammatory response and infiltration of inflammatory cells within aortic tissue is a well-established process in AAA development. Therefore, we aimed to analyze the accompanying inflammation in more details and assessed systemic and vascular inflammation *in vivo* and *ex vivo*. Flow cytometry analysis of peripheral blood revealed a significant increase of the myeloid leukocytes and monocytes in AngII treated *Apoe/Has3*-DKO mice as compared to controls (**Figure 3A-G)**, while lymphocyte numbers were preserved (**Suppl. Figure 4**). However, using ^19^F MRI we revealed that this was surprisingly accompanied by a reduced infiltration of phagocyting immune cells into the aortic wall 7 days after AngII treatment. This finding was further confirmed by flow cytometry also demonstrating decreased numbers of myeloid leukocytes and macrophage numbers in the aortic wall of *Apoe/Has3*-DKO mice (**Figure 4**).

**Figure 3.**
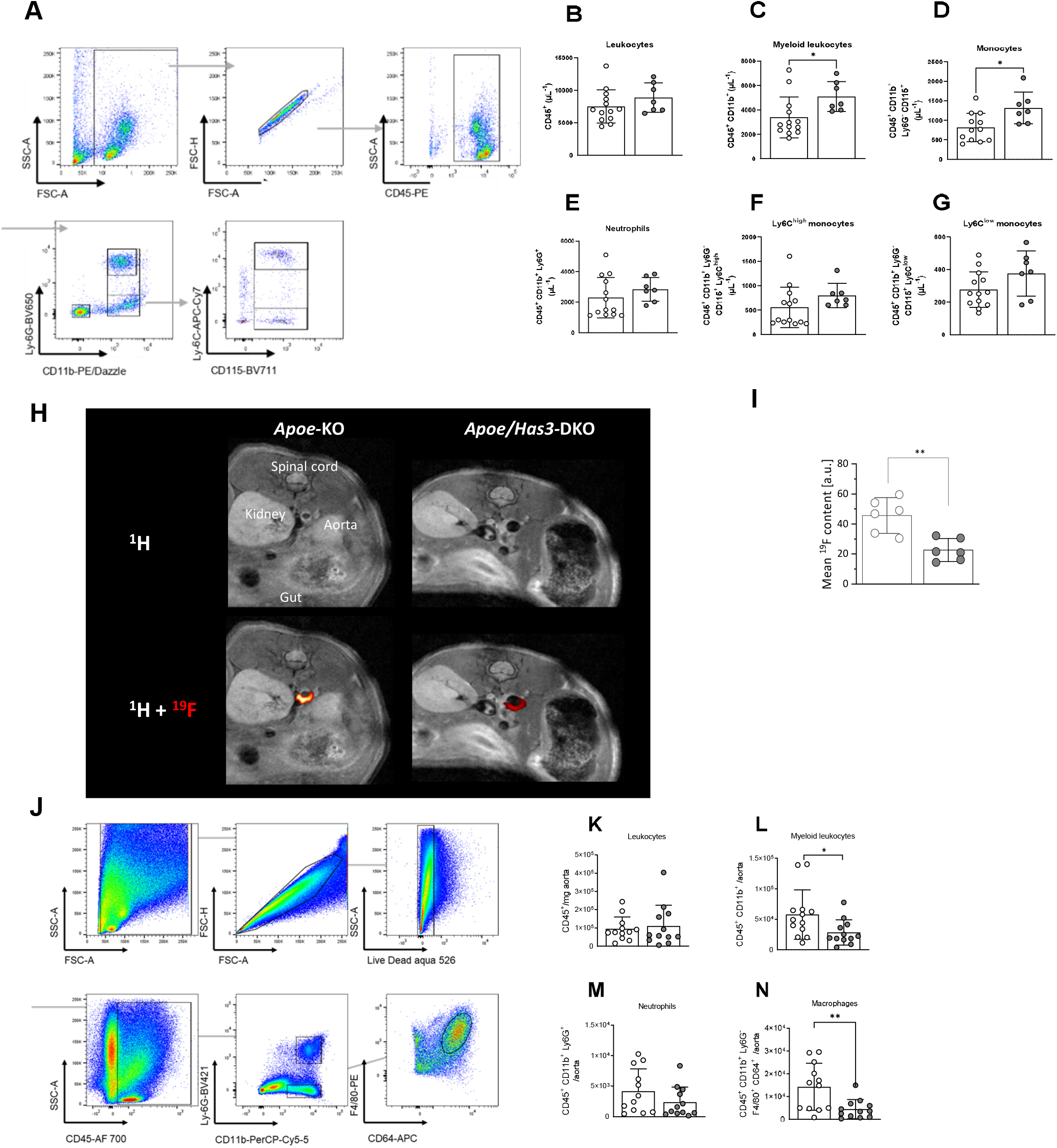
*Has3* deficiency attenuates inflammatory cell infiltration into the aortic wall. **A**, FACS gating strategy to identify inflammatory cell populations in blood. **B-G**, FACS analysis of the peripheral blood revealed a significant increase of **C**, myeloid leukocytes and **D**, monocytes one week after AngII treatment in *Apoe/Ha3*-DKO (n=7) vs. *Apoe*-KO (n=13), * *P* < 0.05, unpaired students *t*-test. **H - I**, ^1^H/^19^F MR inflammation imaging in suprarenal aortas of *Apoe*-KO and *Apoe/Has3*-DKO mice. **H**, Displayed are axial ^1^H scans of the abdominal area (upper panel) and a merging of ^1^H and the aortic ^19^F signal (hot iron scale; lower panel). **I**, Quantification of the ^19^F signal (mean ^19^F signal-to-noise ratio) around the vascular wall (n=6 each group), ** *P* < 0.01, unpaired students *t*-test. **J**, FACS gating strategy to identify inflammatory cell populations in the aortic wall. **K-N**, Fewer **L**, myeloid leukocytes and **N**, macrophages accumulate in the aortic wall of *Apoe/Has3*-DKO (n=12) vs. *Apoe*-KO (n=12), * *P* < 0.05, ** *P* < 0.01, unpaired students *t*-test. **K, M**, *P* ≥ 0.05, unpaired students *t*-test. Data are presented as means ± SD.

**Figure 4.**
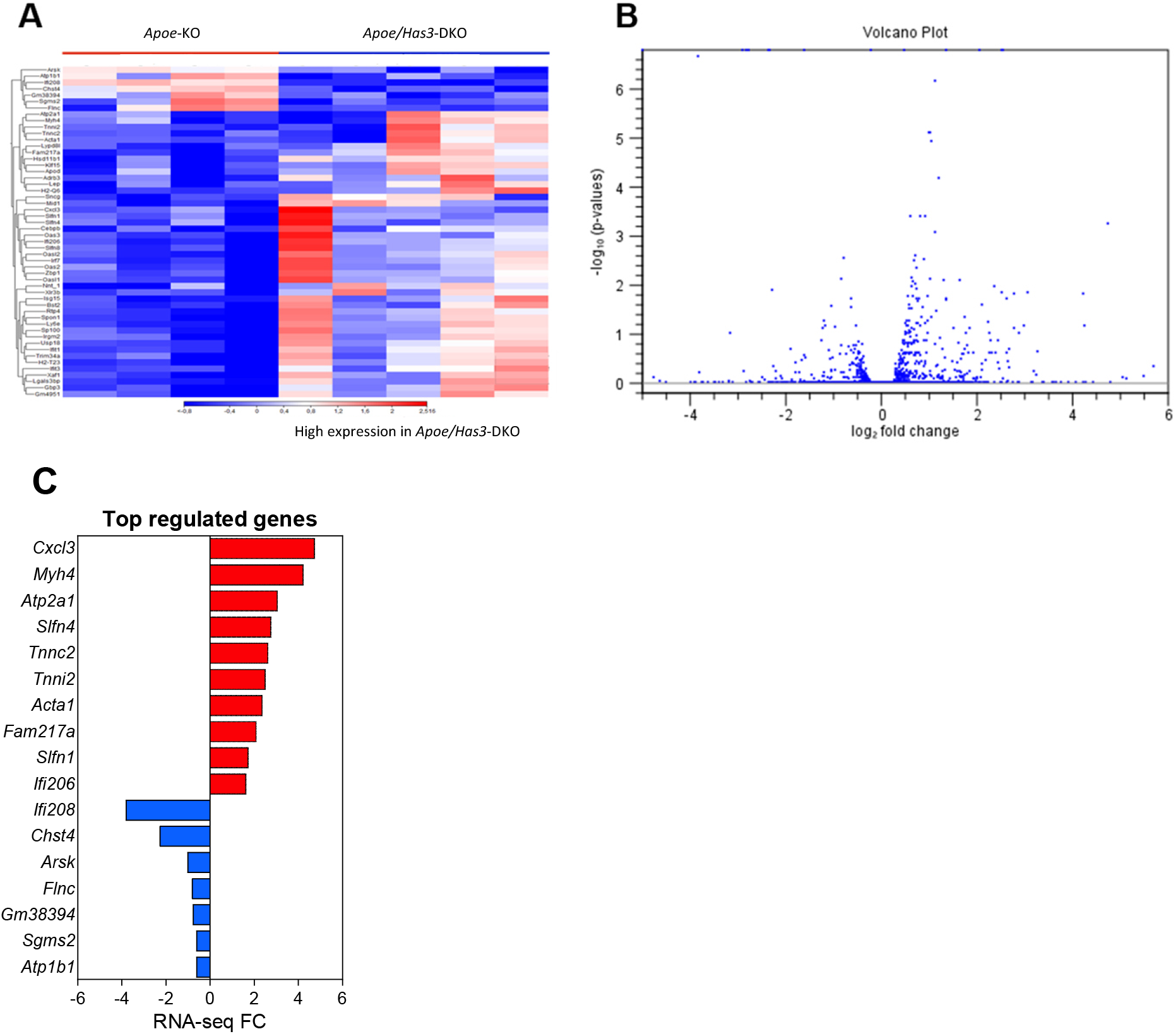
Visualization of differentially expressed genes (DEGs) in aortae of *Apoe/Has3*-KO on day 3 post-AngII infusion identified by bulk RNA-seq assay. **A**, Heatmap of transcriptomic profiling of *Apoe*-KO (n=4) and *Apoe/Has3*-DKO (n=5). Downregulated (blue) and upregulated (red) genes in the aortas of AngII treated *Apoe/Has3*-DKO. **B**, Volcano-plot presenting the differential gene expression between *Apoe/Has3*-DKO and *Apoe*-KO abdominal aortae. Dots represent the product of log_2_-transformed fold change and −log_10_-transformed P values. **C**, Seven genes were downregulated (blue) and 45 were upregulated in the comparison of *Apoe*-KO and *Apoe/Has3*-DKO aortae. Here the 10 most significantly up- and 7 downregulated genes are shown in red and blue accordingly.

Based on these results the underlying mechanism(s) leading to reduced immune cell infiltration into the aortic wall was elucidated. Bulk RNA seq analysis of aortae from *Apoe* and *Apoe/Has3*-DKO mice after three days of AngII treatment revealed 52 differentially expressed genes (DEGs), of which 7 were downregulated and 45 were upregulated in *Apoe/Has3*-DKO mice (**Figure 4CB**). thereby indicating an upregulation of multiple genes and pathways involved in leukocyte adhesion and migration (**Figure 4**). Among the most upregulated markers were *Cxcl3* and *Slfn4* – both important for vascular inflammation and leukocyte infiltration, and GO term analysis revealed “agranulocyte adhesion and diapedesis” as the top regulated pathway (**Table 1**). Additionally, also genes pointing towards a disturbed calcium signaling showed up, which also may contribute to the observed alterations in the aortic phenotype.

**Table 1.** Pathway enrichment analysis for the most statistically significant DEGs in aortas of Apoe/Has3-KO on day 3 post-AngII infusion identified by RNA-seq assay.

| **Name** | **p value** |
| --- | --- |
| Agranulocyte adhesion and diapedesis | $1.81 \times 10^{-10}$ |
| Calcium signaling | $2.63 \times 10^{-8}$ |
| Dilated cardiomyopathy signaling pathway | $5.82 \times 10^{-8}$ |
| Role of pattern recognition receptors in recognition of bacteria and viruses | $9.57 \times 10^{-8}$ |
| Airway pathology in chronic obstructive pulmonary disease | $9.63 \times 10^{-8}$ |

### *Has3* expression in monocytes not endothelial cells regulates aortic infiltration during aortic aneurysm formation

Next the question was addressed whether HAS3 expression in endothelial cells or immune cells is the key determinant for the altered immune cell infiltration into the developing aneurysm. For this, endothelial cells from the abdominal aorta were isolated using a modified Häutchen method (30). However, subsequent bulk RNA sequencing of the isolated endothelial cells revealed no differences between genotypes (**Figure 5A**) thereby excluding endothelial-dependent effects in this process. In contrast, we found mRNA expression of multiple adhesion and diapedesis markers such as *Cx3cr1, Ccr2* and *Ccr5* on isolated monocytes to be upregulated in *Apoe*-KO in response to AngII treatment while no corresponding upregulation was detected in monocytes of *Apoe/Has3*-DKO (**Figure 5B**). Of note, also the HA receptor CD44 was downregulated in *Apoe/Has3*-DKO mice while being upregulated in response to AngII in *Apoe* control mice. Since this receptor has been described to regulate immune-cell-endothelial interactions and promote the adhesion of immune cells to the endothelium, the missing upregulation in response to the pro-inflammatory stimulus is likely to underly the decreased immune cell recruitment in *Apoe/Has3*-DKO mice.

**Figure 5.**
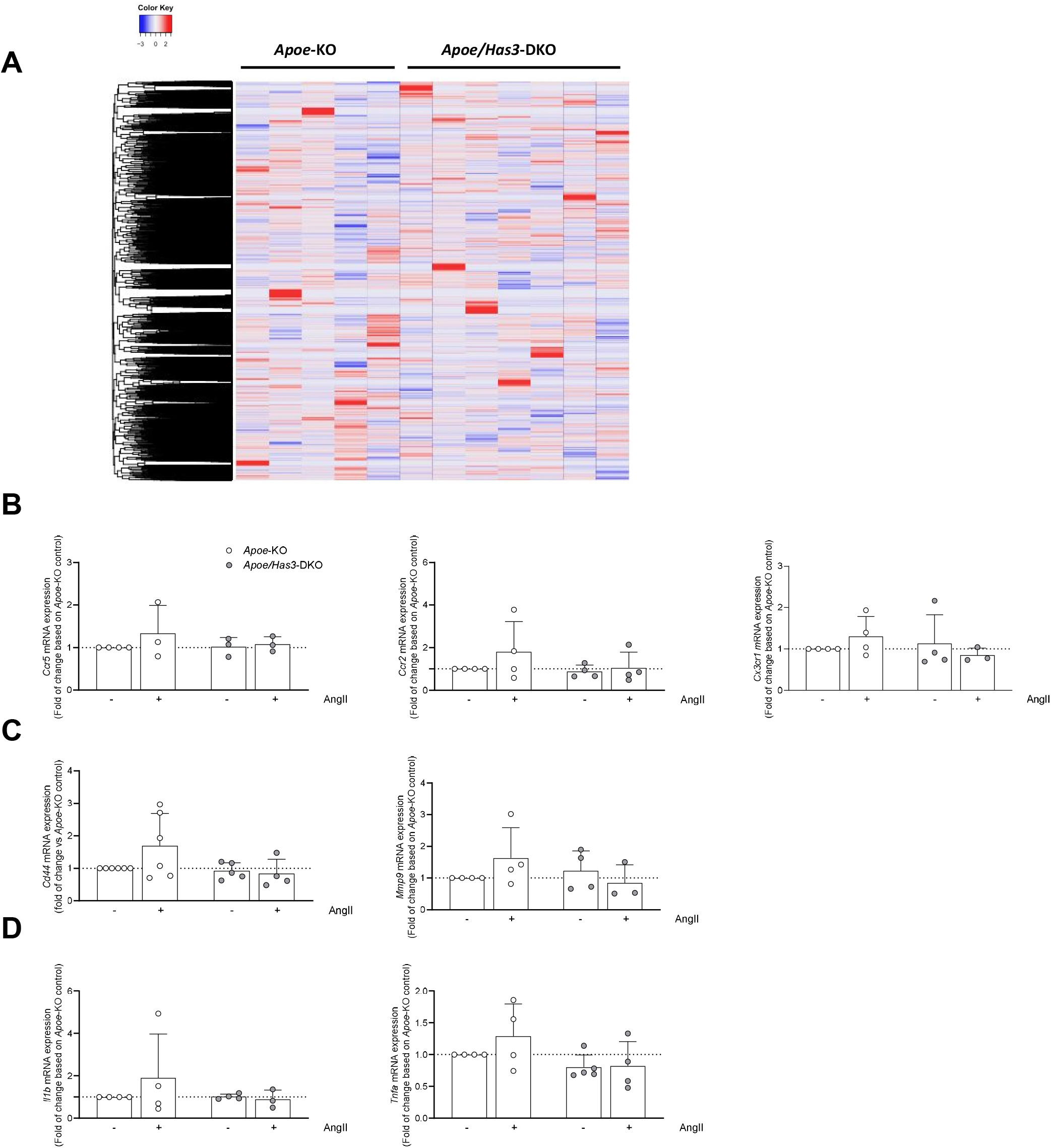
Not endothelial cells but monocytes are responsible for the impaired infiltration into the aortic wall after AngII treatment. **A**, RNAseq heatmap comparing expression of differentially expressed genes in endothelial cells derived from the abdominal aortae of *Apoe*-KO (n=5) and *Apoe/Has3*-KO (n=7) on day 7 post-AngII infusion. In red: upregulation, in blue: downregulation. **B-D**, mRNA expression of **B**, *Ccr5, Ccr2, Cx3cr1*, **C**, *Cd44, Mmp9 und* **D**, *Il1b* and *Tnfa* in primary bone-marrow derived monocytes of *Apoe*-KO (n=3-6) and *Apoe/Has3*-DKO (n=3-5) after 6h of incubation with AngII (10ng/ml) or vehicle. Data are presented as means ± SD.

**Figure 6.**
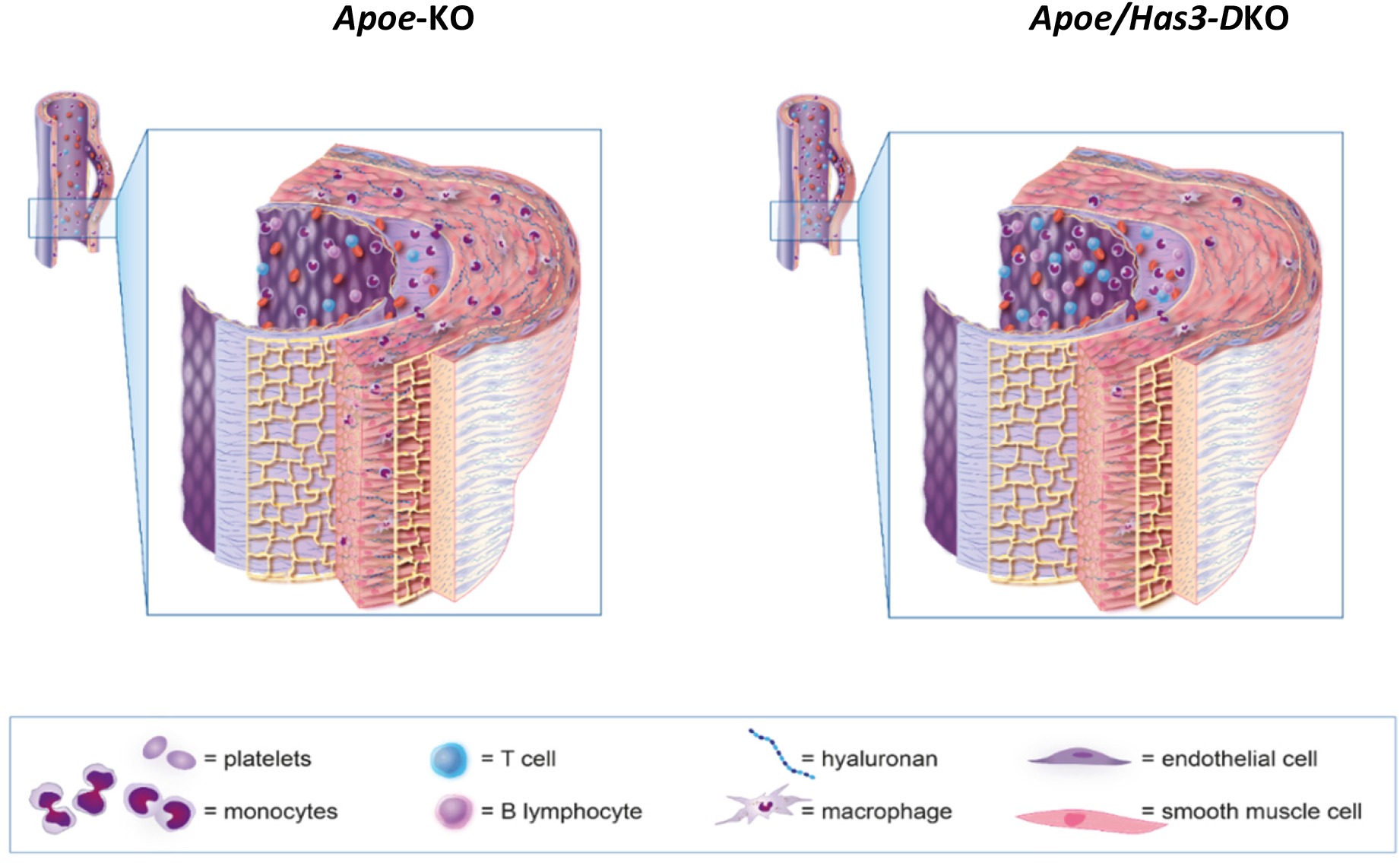
Hyaluronan-rich extracellular matrix and HAS3 promote immune cell infiltration and facilitate ECM degradation in aortic aneurysm and dissection.

## Discussion

In the present study we investigated the role of HAS3-derived HA in the development and progression of aortic aneurysms. In a murine model of Ang-II-induced AAA, we demonstrated that genetic deletion of *Has3* decreased aortic ruptures although aortic diameters were increased. This, at first glance, contradictory findings were based upon reduced numbers of elastica breaks most likely caused by reduced infiltration of leukocytes into the vessel wall. While endothelial cells could be excluded as drivers of this process, monocytes from *Has3*-deficient mice exhibited reduced AngII-mediated upregulation of adhesion markers that are known to be crucial for immune cell adhesion and diapedesis. Most importantly, a strong downregulation of the HA receptor CD44 was observed in monocytes when HAS3 was missing, thereby reducing HA-immune cell interactions and contributing to reduced aortic immune cell infiltration.

Aortic aneurysm formation and progression has been reported to be driven by three major factors: matrix remodeling and degradation (35), inflammatory response(36, 37) as well as smooth muscle cell (SMC) apoptosis. Of note, HA as a major component of the vascular ECM has been described to be involved in all of these processes. While in the present study aortic RNA bulk sequencing gave no evidence for specific alterations in smooth muscle cells, the underlying pathomechanism was instead related to an altered immune cell response. Indeed, previous work highlighted that especially HAS3, one of the three HA synthase isoenzymes, plays a crucial role under a variety of inflammatory conditions such as experimental neointimal hyperplasia (19), experimental colitis (22) or arteriogenesis (21), but in these diseases specifically mediated by effects on vascular SMC (19) or endothelial cells (21, 22). So far, HAS modulation of the inflammatory responses has mainly been observed for T cells, e.g. in atherosclerosis (20) with HAS3-reduced Th1-driven plaque inflammation or myocardial ischemia/reperfusion injury (23) where HAS3 led to less activated and more apoptotic T cells.

Taken together, while these studies clearly support the fundamental role of HA and in particular HAS3-derived HA on immune responses, most of the findings are restricted to T cells. Interestingly, HA-dependent effects have also been reported for macrophages, however, mostly mediated by Toll-like receptors which have been shown to be activated by HA fragments. Here, we report for the first time direct functional effects of HAS3 on monocytes leading to a decreased responsiveness in the used AngII model. Interestingly, an increased adhesion of monocytes under AngII treatment has already been reported in the pathophysiology of hypertension (38, 39). In the present study, we show that lack of HAS3 dampens the Ang-II-induced increase in chemotactic receptor expression. These findings further support the role of HA as a crucial regulator of immune cell invasion. Previous investigation revealed that proinflammatory cytokines such as IL1β and TNFα upregulate HAS2 in endothelial cells thereby promoting monocyte adhesion and recruitment (40). However, here we identified monocyte-derived HA as important regulator for immune cell effector functions under inflammatory conditions as induced by Ang-II. In a previous study, we demonstrated that isolated bone-marrow-derived monocytes not only express all three HAS isoenzymes, but also secrete HA under basal conditions to an even higher amount than differentiated macrophages (29). Therefore, it can be assumed that monocytes produce their own, functional pericellular matrix which also shapes cell functions and in this context monocyte infiltration. This is supported by the finding that mRNA expression of the HA receptor *Cd44* is down-regulated, since CD44 is known to promote immune cell recruitment to the vascular wall (41) and its depletion reduced atherosclerotic lesion size *via* a decreased macrophage recruitment. A similar link might as well represent the underlying mechanism observed in aneurysm formation here. As reported previously for T cells after myocardial ischemia/reperfusion injury (I/R), similarly, in isolated monocytes from mice with AAA *Has3* deficiency was paralleled by decreased *CD44* mRNA expression. One possible mechanism also suggested in the context of I/R might be, that HAS3-derived HA is needed to stabilize CD44 and protect it from proteolytic shedding. However, this relation needs to be addressed in future studies.

Subsequently to reduced monocyte infiltration into the aortic wall, reduced elastica breaks were observed in *Apoe/Has3*-DKO mice compared to *Apoe*-KO littermate controls. The loss of elastic fibres is a key feature during aneurysm development and continously decreases during aneurysm growth (42). In this context, matrix metalloproteinases (MMP) are major drivers of ECM destruction in aortic aneurysm formation (43). Since elastin is preferentially degraded by MMP-2, MMP-9, and MMP-12, they have been extensively studied in AAA. Of note, elevated MMP9 levels can also be detected in patients with AAA. Importantly, monocytes from *Apoe/Has3*-DKO mice not only exhibited decreased AngII-induced expression of adhesion receptors but also showed decreased mRNA expression of *Mmp9*. Although that does not necessarily implies a reduced MMP9 activity, it completes the picture of the underlying mechanisms leading to diminished elastin fragmentation in these mice.

This complex pathomechanism finally results in the reduced rupture incidence in *Apoe/Has3*-DKO and thereby elucidates a novel regulatory role for HAS3 in monocytes during AAA formation.

## Acknowledgement

Computational infrastructure and support were provided by the Centre for Information and Media Technology at Heinrich-Heine-University Düsseldorf. We thank Adrian Brandtner (Institute of Physiology I, University of Bonn) for excellent help with endothelial cell isolation and Julia Odendahl for excellent technical support.

Further we would like to thank Christian Jüngst from the CECAD Imaging Core Facility for adjusting microscopy settings and assistance with recording SHG signals.

## References

1. Sakalihasan N, Michel JB, Katsargyris A, Kuivaniemi H, Defraigne JO, Nchimi A, et al. Abdominal aortic aneurysms. Nat Rev Dis Primers. 2018;4(1):34.

2. Sidloff D, Choke E, Stather P, Bown M, Thompson J, and Sayers R. Mortality from thoracic aortic diseases and associations with cardiovascular risk factors. Circulation. 2014;130(25):2287–94.

3. Han L, Dai L, Zhao YF, Li HY, Liu O, Lan F, et al. CD40L promotes development of acute aortic dissection via induction of inflammation and impairment of endothelial cell function. Aging (Albany NY). 2018;10(3):371–85.

4. Ramanath VS, Oh JK, Sundt TM, 3rd, and Eagle KA. Acute aortic syndromes and thoracic aortic aneurysm. Mayo Clin Proc. 2009;84(5):465–81.

5. An Z, Liu Y, Song ZG, Tang H, Yuan Y, and Xu ZY. Mechanisms of aortic dissection smooth muscle cell phenotype switch. J Thorac Cardiovasc Surg. 2017;154(5):1511–21 e6.

6. Nakashima Y. Pathogenesis of aortic dissection: elastic fiber abnormalities and aortic medial weakness. Ann Vasc Dis. 2010;3(1):28–36.

7. Roberts WC, Vowels TJ, Kitchens BL, Ko JM, Filardo G, Henry AC, et al. Aortic medial elastic fiber loss in acute ascending aortic dissection. The American journal of cardiology. 2011;108(11):1639–44.

8. Duca L, Blaise S, Romier B, Laffargue M, Gayral S, El Btaouri H, et al. Matrix ageing and vascular impacts: focus on elastin fragmentation. Cardiovascular research. 2016;110(3):298–308.

9. Grandoch M, Bollyky PL, and Fischer JW. Hyaluronan: A Master Switch Between Vascular Homeostasis and Inflammation. Circ Res. 2018;122(10):1341–3.

10. Mellak S, Ait-Oufella H, Esposito B, Loyer X, Poirier M, Tedder TF, et al. Angiotensin II mobilizes spleen monocytes to promote the development of abdominal aortic aneurysm in Apoe-/-mice. Arterioscler Thromb Vasc Biol. 2015;35(2):378–88.

11. Raffort J, Lareyre F, Clement M, Hassen-Khodja R, Chinetti G, and Mallat Z. Monocytes and macrophages in abdominal aortic aneurysm. Nature reviews Cardiology. 2017;14(8):457–71.

12. Lu L, Tong Y, Wang W, Hou Y, Dou H, and Liu Z. Characterization and Significance of Monocytes in Acute Stanford Type B Aortic Dissection. J Immunol Res. 2020;2020:9670360.

13. Longo GM, Xiong W, Greiner TC, Zhao Y, Fiotti N, and Baxter BT. Matrix metalloproteinases 2 and 9 work in concert to produce aortic aneurysms. The Journal of clinical investigation. 2002;110(5):625–32.

14. Hu H, Zhang G, Hu H, Liu W, Liu J, Xin S, et al. Interleukin-18 Expression Increases in the Aorta and Plasma of Patients with Acute Aortic Dissection. Mediators Inflamm. 2019;2019:8691294.

15. Choke E, Thompson MM, Dawson J, Wilson WR, Sayed S, Loftus IM, et al. Abdominal aortic aneurysm rupture is associated with increased medial neovascularization and overexpression of proangiogenic cytokines. Arterioscler Thromb Vasc Biol. 2006;26(9):2077–82.

16. Del Porto F, di Gioia C, Tritapepe L, Ferri L, Leopizzi M, Nofroni I, et al. The multitasking role of macrophages in Stanford type A acute aortic dissection. Cardiology. 2014;127(2):123–9.

17. Wen D, Zhou XL, Li JJ, and Hui RT. Biomarkers in aortic dissection. Clin Chim Acta. 2011;412(9-10):688–95.

18. Taylor KR, Yamasaki K, Radek KA, Nardo AD, Goodarzi H, Golenbock D, et al. Recognition of hyaluronan released in sterile injury involves a unique receptor complex dependent on Toll-like receptor 4, CD44, and MD-2. The Journal of biological chemistry. 2007;282(25):18265–75.

19. Kiene LS, Homann S, Suvorava T, Rabausch B, Muller J, Kojda G, et al. Deletion of Hyaluronan Synthase 3 Inhibits Neointimal Hyperplasia in Mice. Arterioscler Thromb Vasc Biol. 2016;36(2):e9–16.

20. Homann S, Grandoch M, Kiene LS, Podsvyadek Y, Feldmann K, Rabausch B, et al. Hyaluronan synthase 3 promotes plaque inflammation and atheroprogression. Matrix biology: journal of the International Society for Matrix Biology. 2018;66:67–80.

21. Schneckmann R, Suvorava T, Hundhausen C, Schuler D, Lorenz C, Freudenberger T, et al. Endothelial Hyaluronan Synthase 3 Augments Postischemic Arteriogenesis Through CD44/eNOS Signaling. Arterioscler Thromb Vasc Biol. 2021;41(10):2551–62.

22. Hundhausen C, Schneckmann R, Ostendorf Y, Rimpler J, von Glinski A, Kohlmorgen C, et al. Endothelial hyaluronan synthase 3 aggravates acute colitis in an experimental model of inflammatory bowel disease. Matrix biology: journal of the International Society for Matrix Biology. 2021;102:20–36.

23. Piroth M, Gorski DJ, Hundhausen C, Petz A, Gorressen S, Semmler D, et al. Hyaluronan synthase 3 is protective after cardiac ischemia-reperfusion by preserving the T cell response. Matrix biology: journal of the International Society for Matrix Biology. 2022;112:116–31.

24. Trachet B, Aslanidou L, Piersigilli A, Fraga-Silva RA, Sordet-Dessimoz J, Villanueva-Perez P, et al. Angiotensin II infusion into ApoE-/-mice: a model for aortic dissection rather than abdominal aortic aneurysm? Cardiovascular research. 2017;113(10):1230–42.

25. Pyo R, Lee JK, Shipley JM, Curci JA, Mao D, Ziporin SJ, et al. Targeted gene disruption of matrix metalloproteinase-9 (gelatinase B) suppresses development of experimental abdominal aortic aneurysms. The Journal of clinical investigation. 2000;105(11):1641–9.

26. Flogel U, Ding Z, Hardung H, Jander S, Reichmann G, Jacoby C, et al. In vivo monitoring of inflammation after cardiac and cerebral ischemia by fluorine magnetic resonance imaging. Circulation. 2008;118(2):140–8.

27. Temme S, Yakoub M, Bouvain P, Yang G, Schrader J, Stegbauer J, et al. Beyond Vessel Diameters: Non-invasive Monitoring of Flow Patterns and Immune Cell Recruitment in Murine Abdominal Aortic Disorders by Multiparametric MRI. Front Cardiovasc Med. 2021;8:750251.

28. Amend SR, Valkenburg KC, and Pienta KJ. Murine Hind Limb Long Bone Dissection and Bone Marrow Isolation. J Vis Exp. 2016(110).

29. Petz A, Grandoch M, Gorski DJ, Abrams M, Piroth M, Schneckmann R, et al. Cardiac Hyaluronan Synthesis Is Critically Involved in the Cardiac Macrophage Response and Promotes Healing After Ischemia Reperfusion Injury. Circ Res. 2019;124(10):1433–47.

30. Simmons CA, Zilberberg J, and Davies PF. A rapid, reliable method to isolate high quality endothelial RNA from small spatially-defined locations. Ann Biomed Eng. 2004;32(10):1453–9.

31. Afgan E, Baker D, Batut B, van den Beek M, Bouvier D, Cech M, et al. The Galaxy platform for accessible, reproducible and collaborative biomedical analyses: 2018 update. Nucleic Acids Res. 2018;46(W1):W537–W44.

32. Dobin A, Davis CA, Schlesinger F, Drenkow J, Zaleski C, Jha S, et al. STAR: ultrafast universal RNA-seq aligner. Bioinformatics. 2013;29(1):15–21.

33. Liao Y, Smyth GK, and Shi W. featureCounts: an efficient general purpose program for assigning sequence reads to genomic features. Bioinformatics. 2014;30(7):923–30.

34. Love MI, Huber W, and Anders S. Moderated estimation of fold change and dispersion for RNA-seq data with DESeq2. Genome Biol. 2014;15(12):550.

35. Sakalihasan N, Heyeres A, Nusgens BV, Limet R, and Lapiere CM. Modifications of the extracellular matrix of aneurysmal abdominal aortas as a function of their size. Eur J Vasc Surg. 1993;7(6):633–7.

36. Newman KM, Jean-Claude J, Li H, Ramey WG, and Tilson MD. Cytokines that activate proteolysis are increased in abdominal aortic aneurysms. Circulation. 1994;90(5 Pt 2):II224–7.

37. Henderson EL, Geng YJ, Sukhova GK, Whittemore AD, Knox J, and Libby P. Death of smooth muscle cells and expression of mediators of apoptosis by T lymphocytes in human abdominal aortic aneurysms. Circulation. 1999;99(1):96–104.

38. Mateo T, Abu Nabah YN, Abu Taha M, Mata M, Cerda-Nicolas M, Proudfoot AE, et al. Angiotensin II-induced mononuclear leukocyte interactions with arteriolar and venular endothelium are mediated by the release of different CC chemokines. J Immunol. 2006;176(9):5577–86.

39. Abu Nabah YN, Losada M, Estelles R, Mateo T, Company C, Piqueras L, et al. CXCR2 blockade impairs angiotensin II-induced CC chemokine synthesis and mononuclear leukocyte infiltration. Arterioscler Thromb Vasc Biol. 2007;27(11):2370–6.

40. Vigetti D, Genasetti A, Karousou E, Viola M, Moretto P, Clerici M, et al. Proinflammatory cytokines induce hyaluronan synthesis and monocyte adhesion in human endothelial cells through hyaluronan synthase 2 (HAS2) and the nuclear factor-kappaB (NF-kappaB) pathway. The Journal of biological chemistry. 2010;285(32):24639–45.

41. Cuff CA, Kothapalli D, Azonobi I, Chun S, Zhang Y, Belkin R, et al. The adhesion receptor CD44 promotes atherosclerosis by mediating inflammatory cell recruitment and vascular cell activation. The Journal of clinical investigation. 2001;108(7):1031–40.

42. Dobrin PB, and Mrkvicka R. Failure of elastin or collagen as possible critical connective tissue alterations underlying aneurysmal dilatation. Cardiovasc Surg. 1994;2(4):484–8.

43. Daugherty A, and Cassis LA. Mechanisms of abdominal aortic aneurysm formation. Curr Atheroscler Rep. 2002;4(3):222–7.

