## Supplement for "Deficiency in hyaluronan synthase 3 attenuates ruptures in a murine model of abdominal aortic aneurysms by reduced aortic monocyte infiltration"

**Suppl. Data**

**Suppl. Table 1. List of the antibodies used for flow cytometry**

| Antibody | Clone | Cat. No. | Vendor | Titer |
|---|---|---|---|---|
| **Blood leukocytes** | | | | |
| CD45-PE | 30-F11 | 103106 | Biolegend | 1:25 |
| Ly6G-BV650 | 1A8 | 127641 | Biolegend | 1:12.5 |
| CD11b-PE/Dazzle | M1/70 | 101256 | Biolegend | 1:100 |
| Ly-6C-APC-Cy7 | HK1.4 | 128026 | Biolegend | 1:200 |
| CD115-BV711 | AFS98 | 135515 | Biolegend | 1:50 |
| **Blood lymphocytes** | | | | |
| CD19-PacBl | 6D5 | 115523 | Biolegend | 1:12.5 |
| CD3-AF700 | 17A2 | 100216 | Biolegend | 1:50 |
| CD8a-AF647 | 53-6.7 | 100724 | Biolegend | 1:100 |
| CD4-FITC | RM4-5 | 11-0042-82 | Thermo Fisher Scientific | 1:200 |
| **Aorta leukocytes** | | | | |
| Live Dead aqua 526 | | L34965 | Thermo Fisher Scientific | 1:25 |
| CD45-AF 700 | 30-F11 | 103128 | Biolegend | 1:25 |
| Ly-6G-BV421 | 1A8 | 127627 | Biolegend | 1:25 |
| CD11b-PerCP-Cy5-5 | M1/70 | 101228 | Biolegend | 1:25 |
| F4/80-PE | BM8 | 123110 | Biolegend | 1:25 |
| CD64-APC | X54-5/7.1 | 139306 | Biolegend | 1:25 |

**Suppl. Table 2. Definitions of immune cell populations quantified by flow cytometry in circulating blood and aorta**

| **Circulating immune cell populations** | **Markers** |
|---|---|
| Myeloid Leukocytes | CD45+CD11b+ |
| Lymphocytes | $CD45^{+}CD19^{+}CD3^{+}$ |
| Monocytes | $CD45^{+}CD11b^{+}Ly6G^{-}CD115^{+}$ |
| Ly6C*high* Monocytes | $CD45^{+}CD11b\ ^{+}CD115^{+}Ly6C^{high}$ |
| Ly6C*low* Monocytes | $CD45^{+}CD11b\ ^{+}CD115^{+}Ly6C^{low}$ |
| Neutrophils | $CD45^{+}CD11b^{+}CD115^{-}Ly6G^{+}$ |
| $CD4^{+}$ T cells | $CD45^{+}CD19^{+}\ Ly6G^{-}CD11b^{-}CD19^{-}CD3^{+}CD4^{+}$ |
| $CD8a^{+}$ T cells | $CD45^{+}CD19^{+}Ly6G^{-}CD11b^{-}CD19^{-}CD3^{+}CD4^{-}CD8a^{+}$ |
| B cells | $CD45^{+}CD19^{+}\ Ly6G^{-}CD11b^{-}CD19^{+}$ |
| **Aortic immune cell populations** | **Markers** |
| Myeloid Leukocytes | CD45+CD11b+ |
| Lymphocytes | $CD45^{+}Ly6G^{-}CD11b^{-}$ |
| Neutrophils | $CD45^{+}CD11b^{+}Ly6G^{+}$ |
| Macrophages | $CD45^{+}CD11b^{+}Ly6G^{-}F4/80^{+}CD64^{+}$ |

**Suppl. Table 3. Murine primer sequences**

| Gene | Forward | Reverse |
|---|---|---|
| *18S* | GCAATTATTCCCCATGAACG | GGCCTCACTAAACCATCCAA |
| *Cd44* | GACCGGTTACCATAACTATTGTC | CATCGATGTCTTCTTGGTGTG |
| *Mmp9* | CCTGAAAACCTCCAACCTCA | GCTTCTCTCCCATCATCTGG |
| *Tnf alpha* | TCGAGTGACAAGCCTGTAGC | AAGGTACAACCCATCGGCTG |
| *Il1 beta* | GGATGAGGACATGAGCACCT | CGTCACACACCAGCAGGTTA |
| *Ccr5* | AGACATCCGTTCCCCCTACA | GCAGGGTGCTGACATACCAT |
| *Ccr2* | AGTTCAGCTGCCTGCAAAGA | GCCGTGGATGAACTGAGGTA |
| *Cx3CR1* | AGTGTGTCGGGTGTCCATTC | GGTAAGGCGAGTCAGCAGTT |

### Suppl. Figure 1

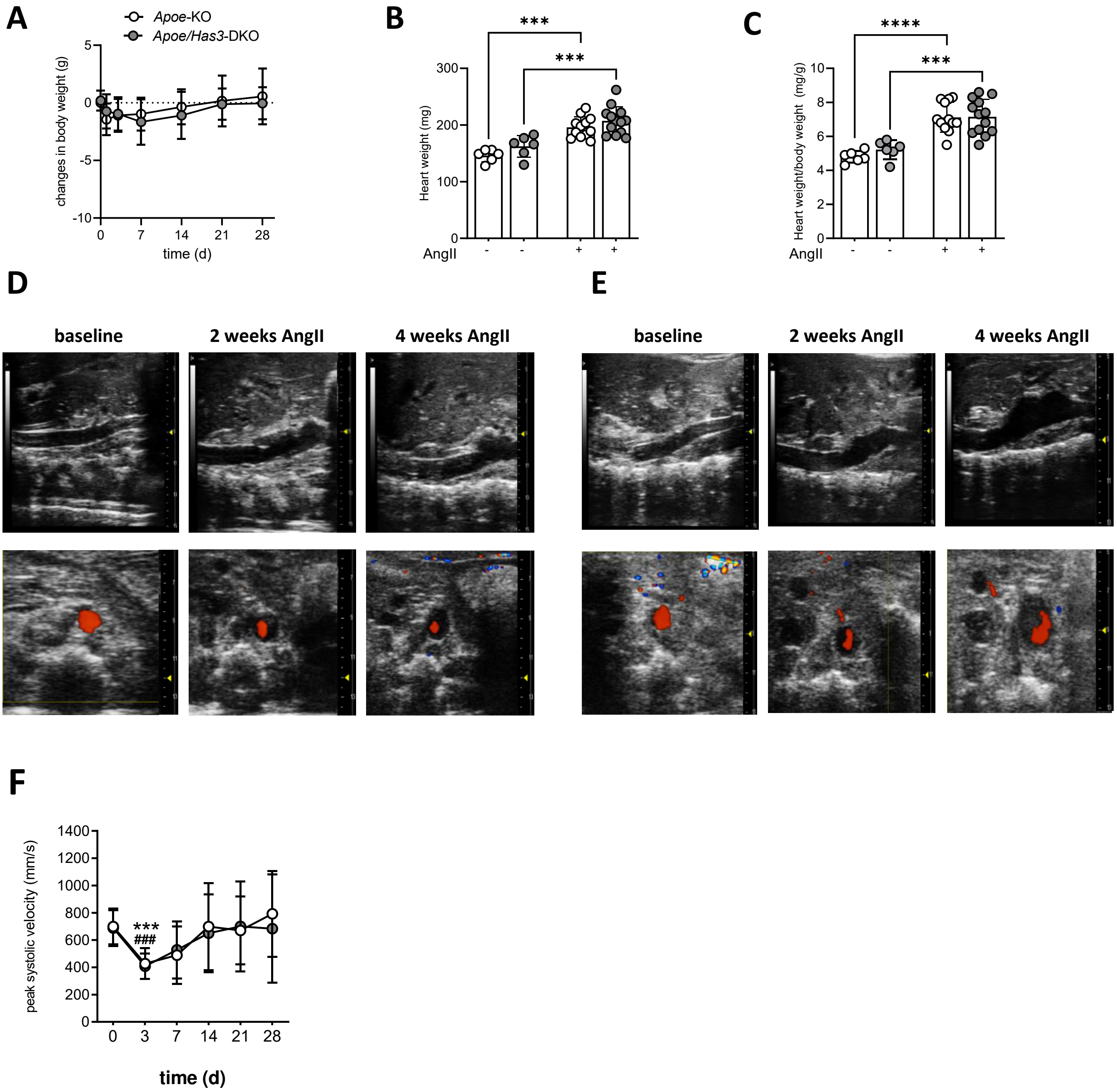


**Suppl. Figure 1. Body weight changes, cardiac hypotrophy and time course of AAA development in *Apoe*-KO and *Apoe/Has3*-DKO mice treated with AngII for 28 days. A,** Body weight changes, Two-way ANOVA, Sidak's post hoc test, **B,** heart weight and **C,** heart weight/body weight ratios in *Apoe*-KO and *Apoe/Has3*-DKO mice, *** $P < 0.001$, unpaired students *t*-test. **D-E,** Representative longitudinal images showing an increase of the aortic diameter (upper panel) and B-mode visualization of the lumen extension and the suprarenal flow with color flow Doppler (red, lower transversepanel) in **D,** *Apoe*-KO and **E,** *Apoe/Has3*-DKO at baseline and on 2 and 4 weeks of AngII infusion. **F,** Blood flow velocity in the suprarenal aorta of *Apoe*-KO (n=11) and *Apoe/Has3*-DKO (n=12) during AngII perfusion, *** $P < 0.001$ and ### $P < 0.0005$ vs. baseline for *Apoe*-KO and *Apoe/Has3*-KO accordingly, Two-way ANOVA, Sidak's post hoc test. Data are presented as means ± SD.

### Suppl. Figure 2

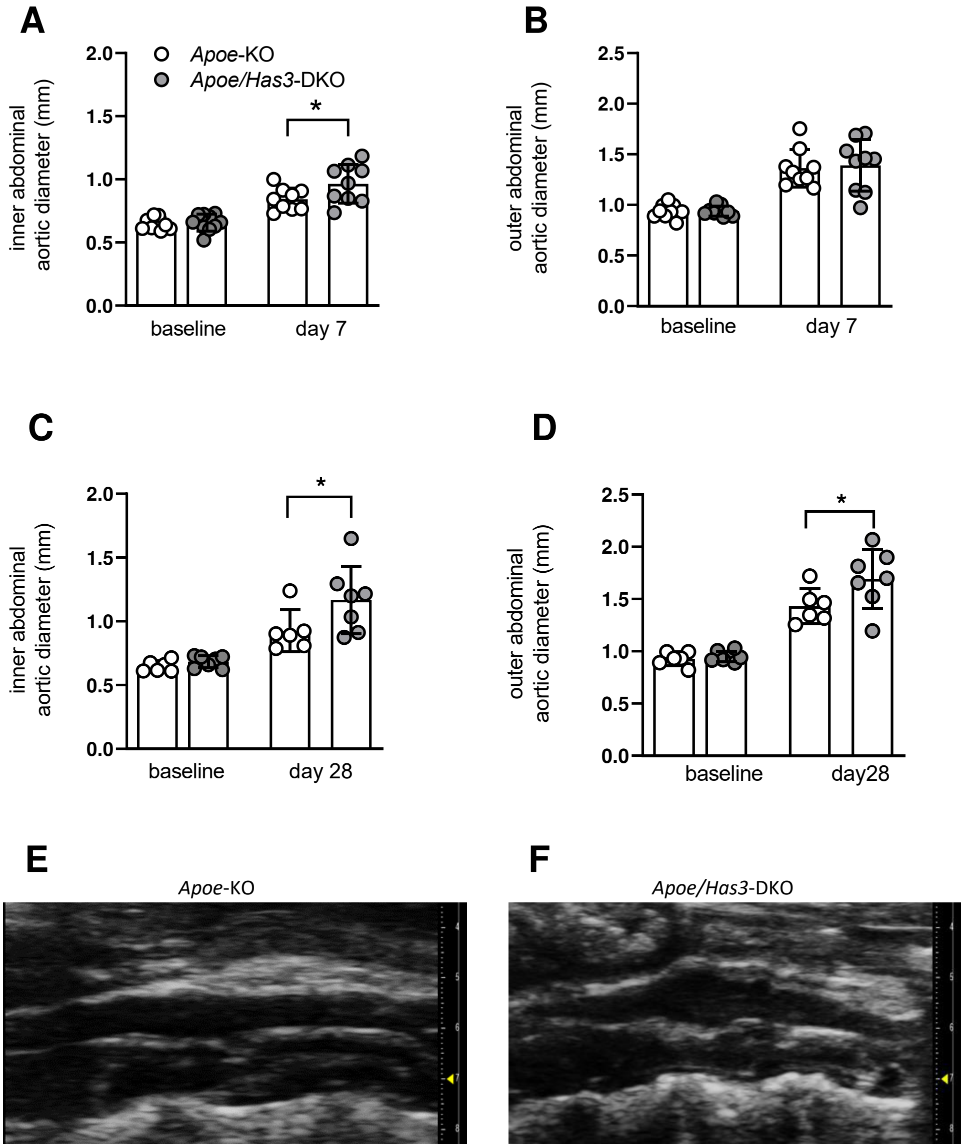


**Suppl. Figure 2.** ***Has3*** **deficiency leads to the augmented infrarenal aneurysm formation in the murine PPE model of AAA. A,C,** Significantly increased inner and **D,** outer aortic diameter after **A, B** day 7 (n=9, each group) and **C,D,** day 28 (n=6, each group) of the PPE perfusion in *Apoe/Has3*-DKO as compared to *Apoe*-KO, * *P* < 0.05, Two-way ANOVA followed by Sidak's post hoc test. **E-F,** Representative longitudinal B-mode images showing an increase of the aortic diameter on day 28 post-PPE perfusion in **E,** *Apoe*-KO and **F,** *Apoe/Has3*-DKO mice. Data are presented as means ± SD.

### Suppl. Figure 3

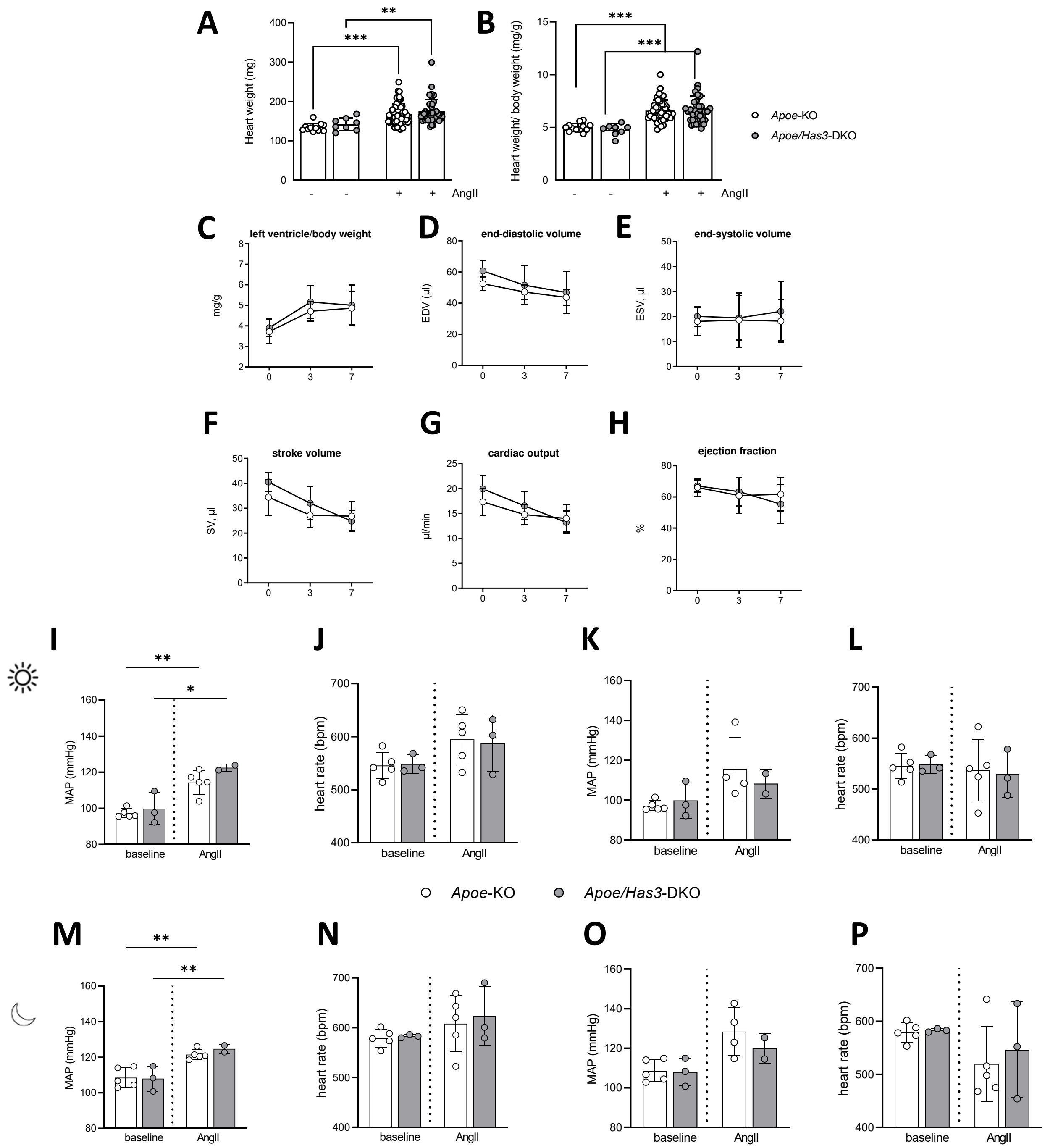


**Suppl. Figure 2. *Has3* deficiency does not affect cardiac function and blood pressure at baseline and in response to one week of AngII infusion. A-B. A,** Heart weight and **B,** heart weight/body weight ratios in *Apoe*-KO and *Apoe/Has3*-DKO mice, ** $P < 0.01$, ***$P < 0.001$, Two-way ANOVA followed by Sidak's post hoc test. **C-H,** Echocardiographic analysis revealed identical **C,** left ventricle hypotrophy and **D-H**, cardiac response to AngII treatment between *Apoe*-KO (n=7) and *Apoe/Has3*-DKO (n=7), **C,** Left ventricular mass/body weight ratio, **D,** end-diastolic volume, **E,** end-systolic volume, **F,** stroke volume, **G,** cardiac output, and **H,** ejection fraction between *Apoe*-KO and *Apoe/Has3*-DKO mice, $P \geq 0.05$, Two-way ANOVA, Sidak's post hoc test. **I-P,** No effect of Has3 deficiency on hemodynamics at baseline and after AngII treatment. **I,K,M,O** Mean arterial blood pressure (MAP) and **J, L, N, P,** heart rate after **I,J,M,N,** 3 days and **K,L,O,P** one week of AngII infusion in *Apoe*-KO (n=5) and *Apoe/Has*3-DKO (n=2-3) mice. **I-L,** diurnal, **M-P**, nocturnal MAP and heart rate ***** $P < 0.05$, ****** $P < 0.005$ vs. baseline, Two-way ANOVA followed by Sidak's post hoc test. Data are presented as means ± SD.

### Suppl. Figure 4

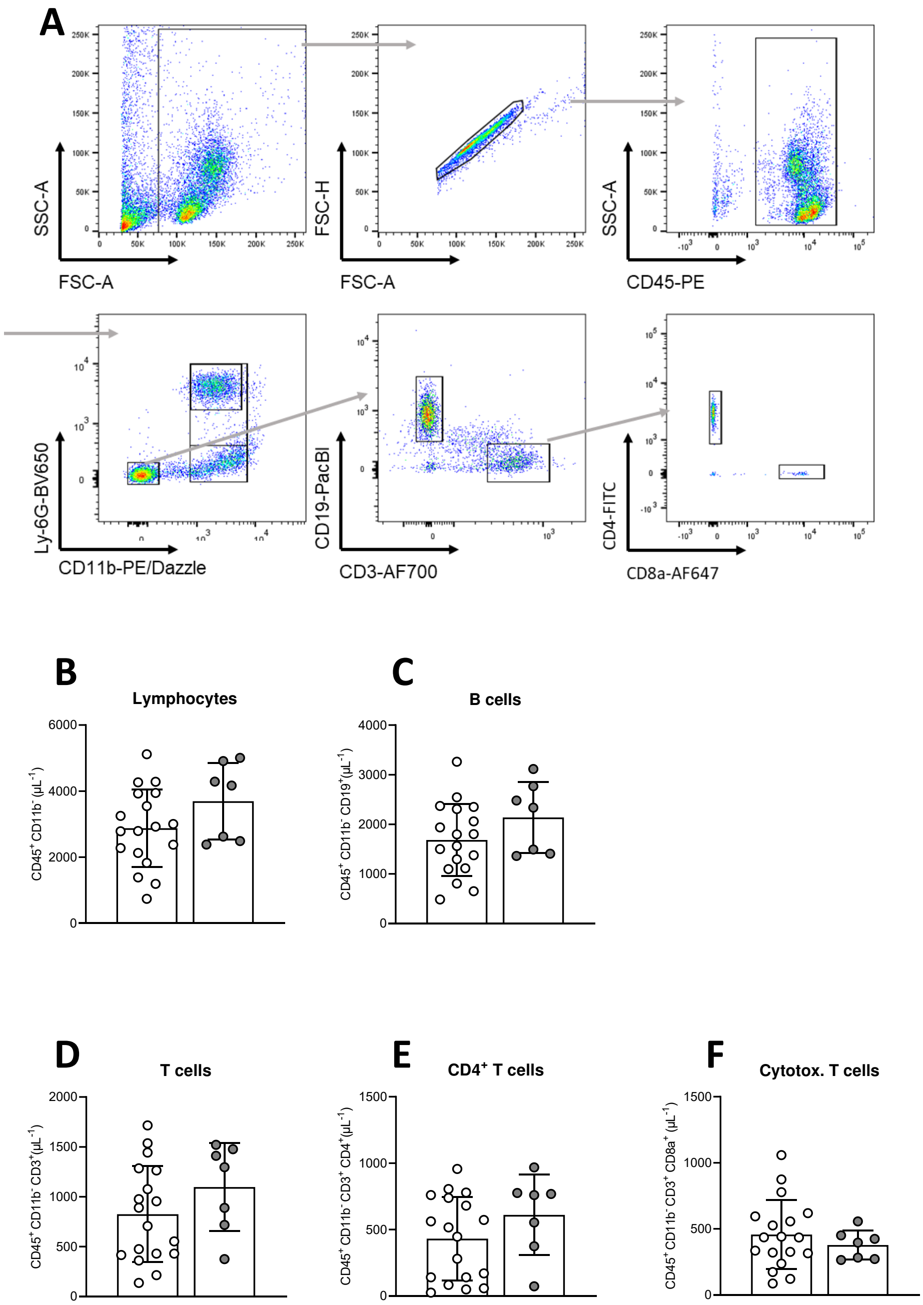


**Suppl. Figure 4. Similar amounts of lymphocytes in peripheral blood of *Apoe*-KO and *Apoe/Has3*-DKO mice on days 7 after AngII perfusion. A**, FACS gating strategy to identify circulating lymphocytes. **B-F,** FACS analysis of the peripheral blood revealed no significant changes in **B,** lymphocytes, **C,** B cells, **D,** T cells, **E,** cytotoxic T cells, and **F,** $CD4^+$ T-cells one week after AngII treatment in *Apoe/Has3*-DKO (n=7) vs. *Apoe*-KO (n=18), P ≥ 0.05, unpaired students *t*-test. Data are presented as means ± SD.
